# Xylanolytic psychrotrophs from andosolic sedge fens and moss heaths in Iceland

**DOI:** 10.1101/348003

**Authors:** Olivia J. Rickman, M. Auður Sigurbjörnsdóttir, Oddur Vilhelmsson

## Abstract

Nine xylanolytic bacterial strains were isolated from fen and heath soils in northern Iceland. They were found by 16S rRNA gene sequencing to belong to the genera *Paenibacillus*, *Bacillus*, *Pseudomonas*, and *Stenotrophomonas*. Using a simple, plate-based semiquantitative assay with azo-crosslinked xylan as the substrate, it was determined that although isolated from cold environments, most of the strains displayed greater xylanolytic activity under mesophilic conditions, with only the paenibacilli displaying markedly cold-active xylanolytic activity. Indeed, for one isolate, *Paenibacillus castaneae* OV2122, xylanolytic activity was only detected at 15°C and below under the conditions tested. Of the nine strains, *Paenibacillus amylolyticus* OV2121 displayed the greatest activity at 5°C. Glycohydrolase family-specific PCR indicated that the paenibacilli produced multiple xylanases of families 10 and 11, whereas a family 8 xylanase was detected in *Pseudomonas kilonensis* AL1515, and a family 11 xylanase in *Stenotrophomonas rhizophila* AL1610.

## Introduction

Xylanases, catalyzing the endohydrolysis of 1,4-β-D-xylosidic linkages in the hemocellulosic polymer xylan, are a widespread group of enzymes that play an important role in plant detritus degradation and are of considerable importance in many industrial processes, such as in the paper and food industries (3, 7). Unsurprisingly, xylanase-producing microorganisms are typically found on plants or in association with plant material, such as in the phyllosphere and rhizosphere (15, 32). While xylanases have been isolated from various extremophiles, in particular thermophiles (7), research on cold-adapted xylanases is relatively scarce. Several xylanases have nevertheless been isolated from psychrophiles (12, 19) and found to possess common features such as a low temperature optimum and poor thermostability due to fewer salt-bridges and less compact hydrophobic packing as compared to their mesophilic and thermophilic counterparts (7). Xylanolytic activity is among the most important polymer degradation activities in the phyllosphere and xylanolytic bacteria can thus be expected to be important players in plant detritus turnover. Furthermore, xylanases are valuable enzymes that are used in several processes within the food and paper industries. Cold active xylanases and psychrotrophic xylanolytic organisms are thus of considerable interest both to biotechnology and cold-climate ecology.

In this study, we present nine psychrotrophic xylanolytic bacterial strains from three soil habitat types in northern Iceland, identified as paenibacilli, pseudomonads, a *Bacillus* and a *Stenotrophomonas*, and discuss the temperature-dependence of their xylanase activity and the likely glycohydrolase family of their respective xylanases.

## Materials and methods

### Isolation of bacteria

Leptosolic and andosolic sites (Table 1) were sampled aseptically with sterile implements into sterile Falcon tubes (Becton-Dickinson, Franklin Lakes, NJ, USA) or WhirlPak bags (Nasco, Fort Atkinson, WI, USA). To obtain microbial suspensions, soil samples were placed in a tenfold volume of Butterfield’s buffer (Butterfield 1932) at pH 7.2 in a sterile stomacher bag and stomached in a LabBlender 400 (Seward, Worthing, UK) for 2 min. The slurry was then serially diluted in Butterfield’s buffer to 10^−6^. All dilutions were spread-plated in duplicate onto Plate Count Agar (PCA), Tryptic Soy Agar (TSA), and Actinomycete Isolation Agar (AIA) (Becton-Dickinson) and plates incubated in the dark at 15°C for four weeks. The plates were examined for colony morphotypes based on color, sheen, convexity and other visible features. Representatives of each morphotype were aseptically picked, streaked onto fresh media and incubated under identical conditions as the original isolation plates. In order to obtain pure cultures, the isolates were re-streaked and incubated at least once more. Stocks of purified isolates were prepared by suspending a loopful of growth in 0.5 ml 28% (v/v) glycerol and stored at −70 °C.

**Table 1.** Strains, source samples, and isolation media; Plate Count Agar (PCA), Tryptic Soy Agar (TSA) and Actinomycete Isolation Agar (AIA).

| Depth<br>(cm) | Strain no. | Isolation<br>medium | Colony morphology |
| --- | --- | --- | --- |
| **Site 1. Sandabotnaskarð** (65.685°N, 16.769°W). |  |  |  |
| Brown andosol. Sparse vegetation. |  |  |  |
| 10 | AIB0216B | PCA | White, flat, dry |
| 10 | AIB0216C | PCA | White, flat, dry |
| 10 | AIB0228 | TSA | Cream-colored, convex |
| **Site 2. Svartárkot** (65.336°N, 17.242°W) |  |  |  |
| Brown and histic andosol. Moist moss heath. |  |  |  |
| 5 | AL1515 | PCA | Cream-colored, convex |
| 17 | AL1610 | AIA | Tan, convex, glistening, small |
| 17 | AL1614 | PCA | White, matte, convex, grainy |
| 25 | DO1702 | AIA | Cream-colored, irregular, glistening, slimy |
| **Site 3. Hesteyrarskarð** (66.358°N, 22.930°W) |  |  |  |
| Histic andosol. Short sedge fen. |  |  |  |
| 5 | OV2120 | TSA | White, flat, dry |
| 5 | OV2121 | TSA | White, flat, dry |
| 5 | OV2122 | TSA | White, flat, dry |

### Xylan degradation screens and assays

Degradation of xylan was screened for by the appearance of blue halos on an 0.5 g l^−1^ suspension of azo-cross-linked birch xylan (Megazyme,Wicklow, Ireland) in a medium that consisted of 4 g Nutrient Broth (NB) and 15 g l^−1^ agar (Becton-Dickinson). When used as a semi-quantitative xylanase assay, halo emergence on equal-volume (20 ml) plates was monitored twice daily and halo diameter measured with a ruler. One-way ANOVA on terminal halo diameters was used to determine effect of temperature on xylan degradation. Precultures were incubated in the presence of xylan.

### Phenotypic characterization

Several standardized biochemical tests were performed using the API 20E test strips (Biomériaux) according to the manufacturer‘s instructions.

### 16S rRNA gene sequencing

Colonies were picked using sterile toothpicks and suspended in 25 μl colony lysis buffer (1% Triton X-100 in 0.02 mol l^−1^ Tris / 0.002 mol l^−1^ EDTA at pH 8.0). The resulting suspension was heated to 95°C for 10 min and then cooled to 4°C and 1 μl of the suspension used as a template in a standard Taq-PCR reaction (35 cycles, annealing at 51°C for 30 sec, extension at 68°C for 90 sec, denaturing at 95°C for 30 sec) in an MJR PTC-200 thermocycler (MJ Research Inc., Waltham, MA, USA). The primers 8F (5‘-AGAGTTTGATCCTGGCTCAG-3’) and 1522R (5‘-AAGGAGGTGATCCAACCGCA-3’) were used at a final concentration of 0.2 μM in a total volume of 25 μl of PCR mixture containing 0.15 μl of Taq DNA polymerase (New England Biolabs, Ipswich, MA, USA). Amplicons were visualized on an 0.8% agarose gel using SYBR Safe (Life Technologies, Carlsbad, CA, USA) and cleaned up for sequencing using 20 U μl^−l^ exonuclease I and 5 U μl^−1^ Antarctic phosphatase (New England Biolabs) at 37°C for 30 min, followed by deactivation at 90°C for 5 min. Partial sequencing of the purified amplicons was performed with a BigDye terminator kit and run on Applied Biosystems 3130XL DNA analyzer (Applied Biosystems, Foster City, USA) at Macrogen Europe, Amsterdam, the Netherlands using sequencing primers 519F (5‘-CAGCAGCCGCGGTAATAC-3’) and 926R (5‘-CCGTCAATTCCTTTGAGTTT-3’). Sequences were examined in ABI Sequence Scanner 1.0 (Applied Biosystems, Framingham, MA, USA) and the forward and reverse complement of the reverse sequence manually aligned and merged.

### Phylogenetic analysis

The 16S rRNA gene sequences were assigned to taxa based on the identification of phylogenetic neighbors as determined by a BLASTN (1) search against a database containing type strains with validly published prokaryotic names and representatives of uncultured phylotypes (18). The top thirty sequences with the highest scores were then selected for the calculation of pairwise sequence similarity using a global alignment algorithm (22), which was implemented at the EzTaxon server (18). For further phylogenetic analysis, sequences were multiply aligned using MUSCLE (9) and bootstrapped (10) neighbor-joining trees (33) were calculated in MEGA6 (36) using the Maximum Composite Likelihood model (35).

### Amplification of xylanase gene fragments

Genomic DNA was extracted and purified from cultures grown on Nutrient Agar (NA) plates using MoBio Microbial DNA Isolation kit (MoBio Laboratories, Solana Beach, CA) according to the manufacturer’s instructions. Primers specific to xylanases of families 8 (Xyn8A, forward: 5‘-CATCCTGTTCAGGAAGACAGTAGTGGGG-3’, reverse: 5‘-CTTGATAATCCGGAAATTGCCACTGACATGC-3’), 10 (XynFA, forward: 5‘-CACACKCTKGTKTGGCA-3’, reverse: 5‘-TMGTTKACMACRTCCCA-3’), and 11 (PAXynA, forward: 5‘-GAYTAYTGGCARTAYTGGAC-3’, reverse: 5‘-ATRTCRTANGTNCCNCCRTCRCT-3’) were used for amplification. Reaction conditions, amplicon clean-up and sequencing were as described for the 16S rRNA gene amplification, above.

## Results

### Identification of xylanolytic isolates

Isolated bacterial strains (106) from andosolic wetland and moss heath sites (Table 1) were screened for the ability to yield blue halos on azo cross-linked xylan plates at 15°C, and nine positive isolates selected for further study. The isolates were identified by 16S rRNA gene sequencing (Table 2) and found to comprise paenibacilli (5 isolates), pseudomonads (2 isolates), a stenotrophomonad and a *Bacillus*.

**Table 2.** Identification of xylanolytic bacterial isolates by 16S rRNA gene sequencing.

| Isolate | Sequence length (nt) | GenBank accession | Most similar taxon at EzTaxon<sup>†</sup> (%) |
| --- | --- | --- | --- |
| AIB0216B | 1523 | KX349197 | *Paenibacillus tundrae* (99,87) |
| AIB0216C | 1452 | KX349198 | *Paenibacillus tundrae* (99,79) |
| AIB0228 | 1436 | KX349200 | *Pseudomonas baetica* (99.30) |
| AL1515 | 998 | KX349201 | *Pseudomonas kilonensis* (99.50) |
| AL1610 | 1503 | KX349202 | *Stenotrophomonas rhizophila* (99.73) |
| AL1614 | 1435 | KX349199 | *Bacillus subtilis* (99.23) |
| OV2120 | 1459 | KX349195 | *Paenibacillus amylolyticus* (99,72) |
| OV2121 | 1451 | KX349194 | *Paenibacillus amylolyticus* (99,79) |
| OV2122 | 1457 | KX349196 | *Paenibacillus castaneae* (99,73) |

<sup>†</sup> The isolates were identified using the EzTaxon server (http://www.ezbiocloud.net/eztaxon;
Kim et al., 2012) on the basis of 16S rRNA gene sequence data.

A pairwise distance comparison of the MUSCLE multiply aligned 16S rRNA gene sequences obtained for all nine strains (Table 3) reveals that four of the paenibacilli, strains OV2120, OV2121, AIB0216B, and AIB0216C are very closely related to one another, with on average less than 2 base substitutions over the 987 nucleotide positions tested.

**Table 3.**
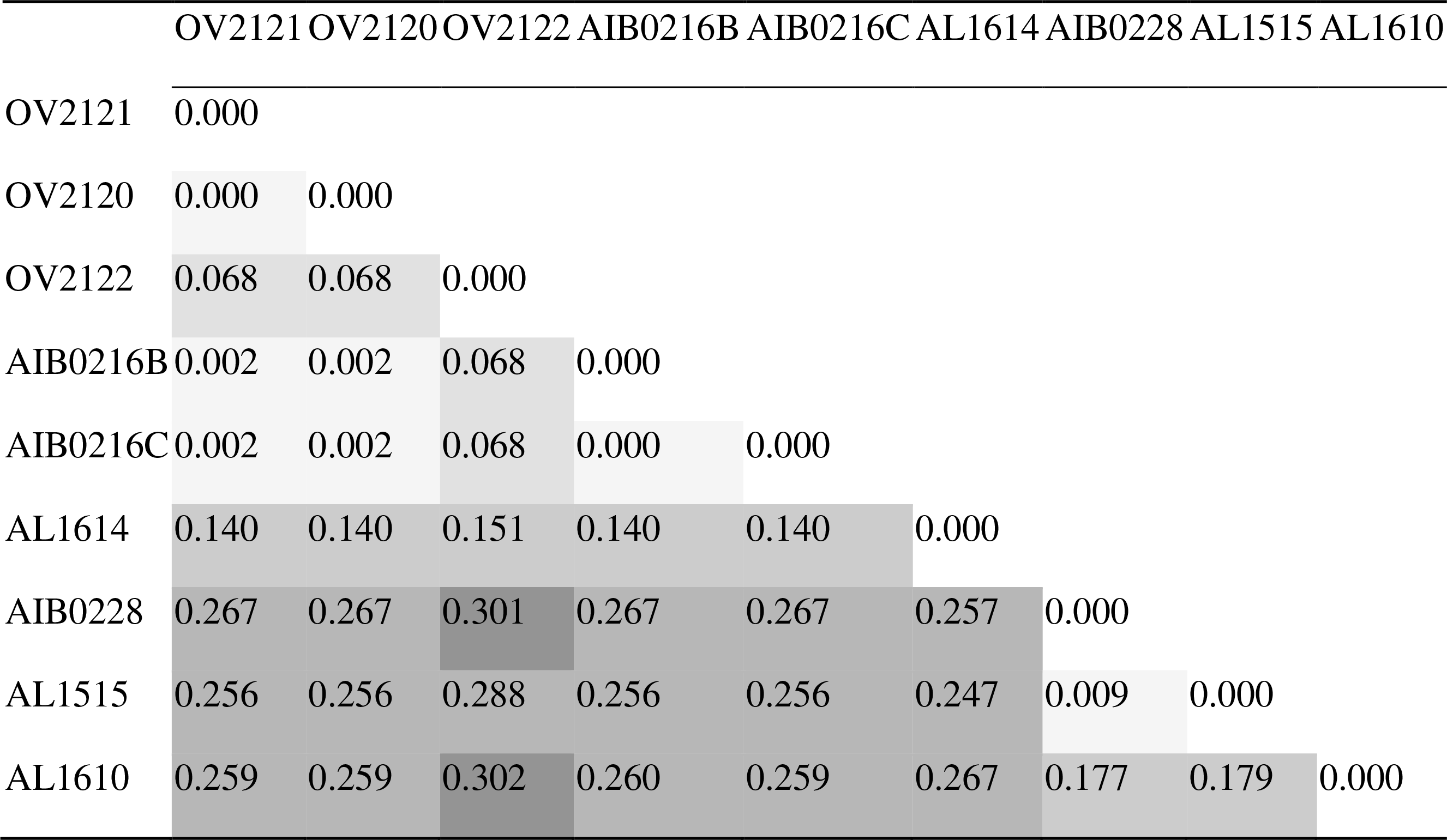
Pairwise distances between MUSCLE multiply aligned 16S rRNA gene sequences.

|  | OV2121 | OV2120 | OV2122 | AIB0216B | AIB0216C | AL1614 | AIB0228 | AL1515 | AL1610 |
| --- | --- | --- | --- | --- | --- | --- | --- | --- | --- |
| OV2121 | 0.000 |  |  |  |  |  |  |  |  |
| OV2120 | 0.000 | 0.000 |  |  |  |  |  |  |  |
| OV2122 | 0.068 | 0.068 | 0.000 |  |  |  |  |  |  |
| AIB0216B | 0.002 | 0.002 | 0.068 | 0.000 |  |  |  |  |  |
| AIB0216C | 0.002 | 0.002 | 0.068 | 0.000 | 0.000 |  |  |  |  |
| AL1614 | 0.140 | 0.140 | 0.151 | 0.140 | 0.140 | 0.000 |  |  |  |
| AIB0228 | 0.267 | 0.267 | 0.301 | 0.267 | 0.267 | 0.257 | 0.000 |  |  |
| AL1515 | 0.256 | 0.256 | 0.288 | 0.256 | 0.256 | 0.247 | 0.009 | 0.000 |  |
| AL1610 | 0.259 | 0.259 | 0.302 | 0.260 | 0.259 | 0.267 | 0.177 | 0.179 | 0.000 |

A phylogenetic analysis of the paenibacilli (Figure 1) was performed using 54 *Paenibacillus* type strain reference sequences. The reference sequences were selected based firstly on their similarity to the test strains as determined by the BLASTN algorithm on the EzTaxon server and secondly based on their completeness, enabling us to run the phylogenetic analysis over 1369 nucleotide positions. The analysis revealed that isolates OV2121, OV2120, AIB0216B, and AIB0216C form a clade with *P. amylolyticus*, *P. xylanexedens*, *P. tundrae* and *P. tylopili*, supporting the EzTaxon assignments of isolates OV2120 and OV2121 as *P. amylolyticus*, and AIB0216B and AIB0216C as *P. tundrae*. The divergence of strain OV2122 from the other four paenibacilli was further demonstrated on the phenotypic level by biochemical characterization using the API 20E strip test, in which OV2122 was distinguished by lack of ONPG hydrolysis, presence of arginine dihydrolase and gelatinase, citrate utilization, and lack of sucrose and amygdalin utilization (Table 4).

**Table 4.**
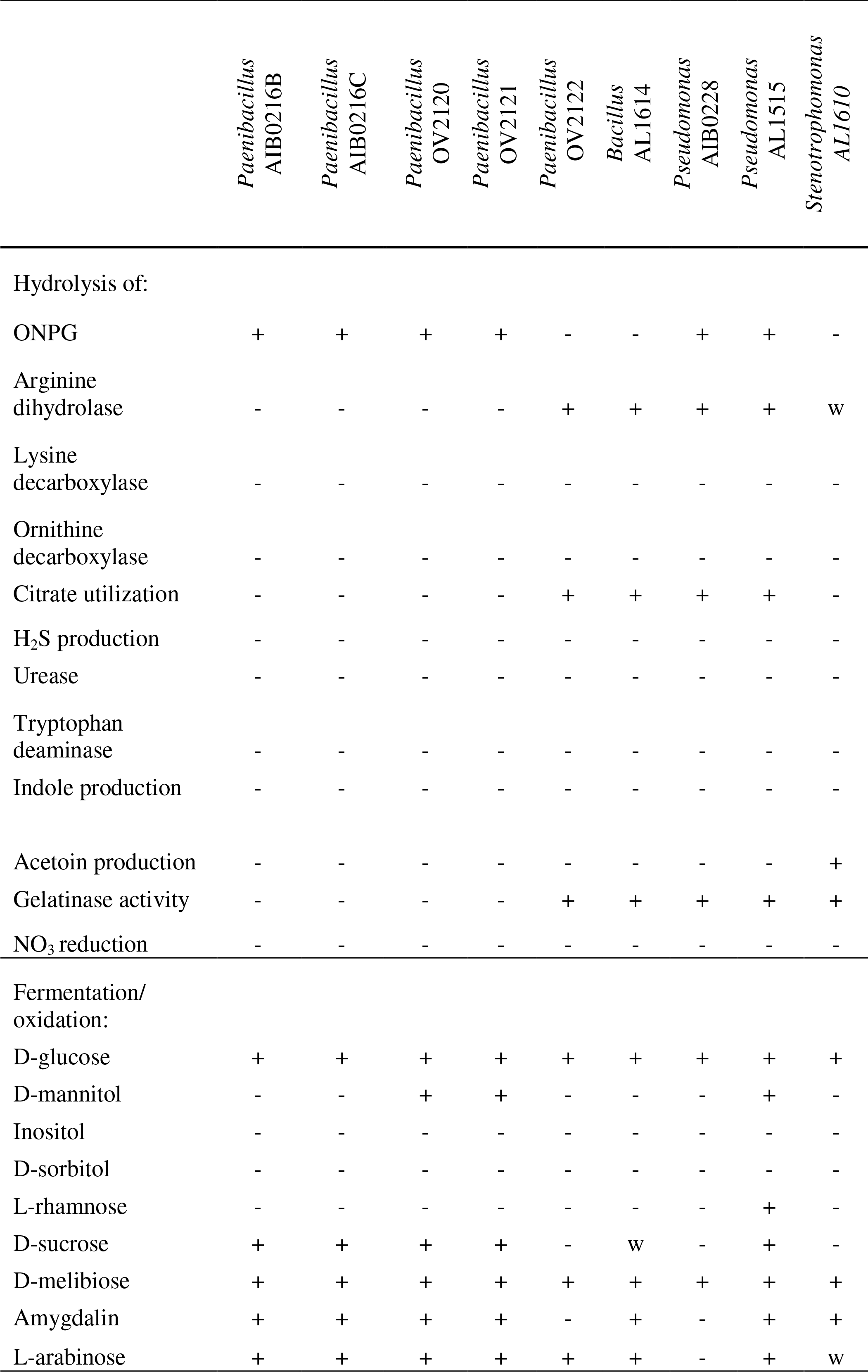
Selected phenotypic characteristics (positive (+), negative (-) or weak (w)) of xylanolytic isolates obtained using API 20E test strips

**Figure 1.**
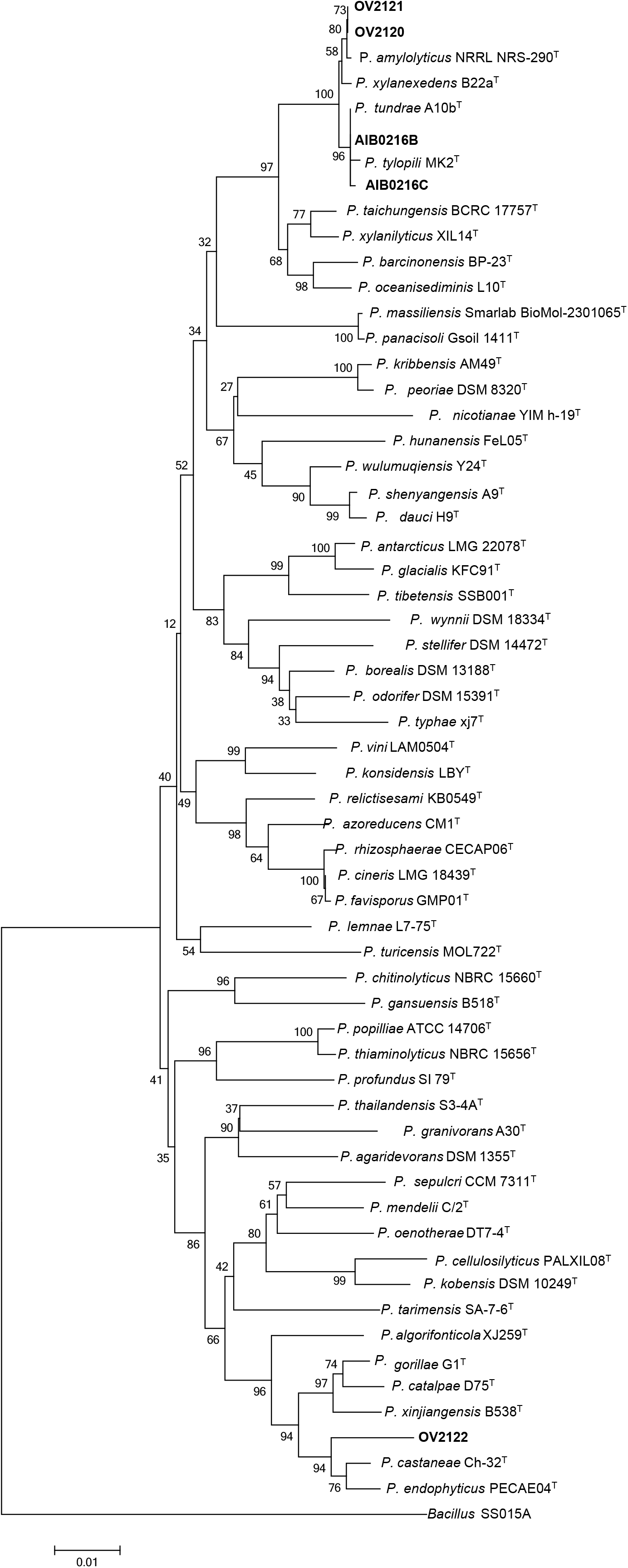
16S rRNA gene phylogenetic analysis of the *Paenibacillus* isolates and selected type strains from that genus. Phylogenetic relationships were determined using the Neighbor-Joining method of Saitou and Nei (33). The optimal tree had a branch length sum of 0.8577. The percentage of replicate trees in which the associated taxa clustered together in the bootstrap test (1000 replicates) are shown next to the branches (10). The tree is drawn to scale, with branch lengths in the same units as those of the evolutionary distances used to infer the phylogenetic tree. The evolutionary distances (number of base substitutions per site) were computed using the Maximum Composite Likelihood method of Tamura et al. (35). All positions containing gaps and missing data were eliminated, resulting in a total of 1369 positions in the final dataset. The analysis was conducted in MEGA6 (36).

### Presence and family of xylanase genes

To independently verify the presence of glycohydrolase family 8, 10 or 11 xylanase genes, PCR reactions using primer pairs Xyn8A, XynFA, and PAXynA were run on DNA extracted from pure cultures of isolated strains, as well as of the Az-Xylan screen-negative strain DO1702 as a negative control (Figure 2). Selected product bands were extracted and sequenced (Table 5). Under the reaction conditions employed, the family 8 primer pair Xyn8A yielded apparent primer dimers in all cases, but nevertheless clear product bands at approximately 400 bp were visible for strains AL1515, OV2121, and OV2122. The AL1515 band was sequenced and found by BLASTx to contain a 359-bp product most similar to a hypothetical protein of unknown function in the Betaproteobacterium *Andreprevotia chitinilytica*, and thus probably constitutes an unspecific amplification of an unknown component of the AL1515 genome, or possibly a hitherto unidentified xylanase. The family 10 primer pair XynFA yielded a variety of amplicons: a large (> 10 kb) product from strain AL1610, an approximately 150 bp product from strain OV2122, an approximately 250 bp product from strain OV2122, three products (3 kb, 1 kb and 700 bp) from each of strains OV2120 and OV2121, and two products (150 and 750 bp) from strain AIB0216B. Of these, three amplicons were sequenced (both AIB0216B products and the AIB0228 product), one of which was found to correspond to a *Paenibacillus* xylanase, while the others showed greatest similarity to proteins of unknown function (Table 5). The family 11 primer pair PAXynA yielded an approximately 200 bp-product from strain AIB0216C, and approximately 800-bp products from strains AL1610, OV2120, OV2121, and OV2122. The products for AL1610, OV2121, and OV2122 were partially sequenced and all displayed greatest similarity to *Paenibacillus* endo-1,4-beta xylanases (Table 5).

**Table 5.** BLASTX of partial nucleotide sequences obtained from amplicons obtained using xylanase-specific PCR primers.

| band | primer | lgth<br>(nt) | best BLASTx hit | %id(aa) |
| --- | --- | --- | --- | --- |
| AL1610 | PAXynA | 261 | *Paenibacillus* sp. HY8 endo-xylanase | 99 |
| OV2122 | PAXynA | 263 | *Paenibacillus amylolyticus* 1,4-beta-xylanase | 99 |
| OV2121 | PAXynA | 204 | *Paenibacillus* sp. PAMC 26794 endo-1,4-beta-xylanase | 95 |
| AIB0216B_b1 | XynFA | 157 | *Paenibacillus* sp. O199 1,4-beta-xylanase | 100 |
| AIB0216B_b2 | XynFA | 768 | *Paenibacillus* sp. O199 hypothetical protein | 100 |
| AIB0228 | XynFA | 247 | *Pseudomonas* sp. RIT-PI-r LOG family protein | 100 |
| AL1515 | Xyn8A | 359 | *Andreprevotia chitinilytica* hypothetical protein | 32 |

**Figure 2.**
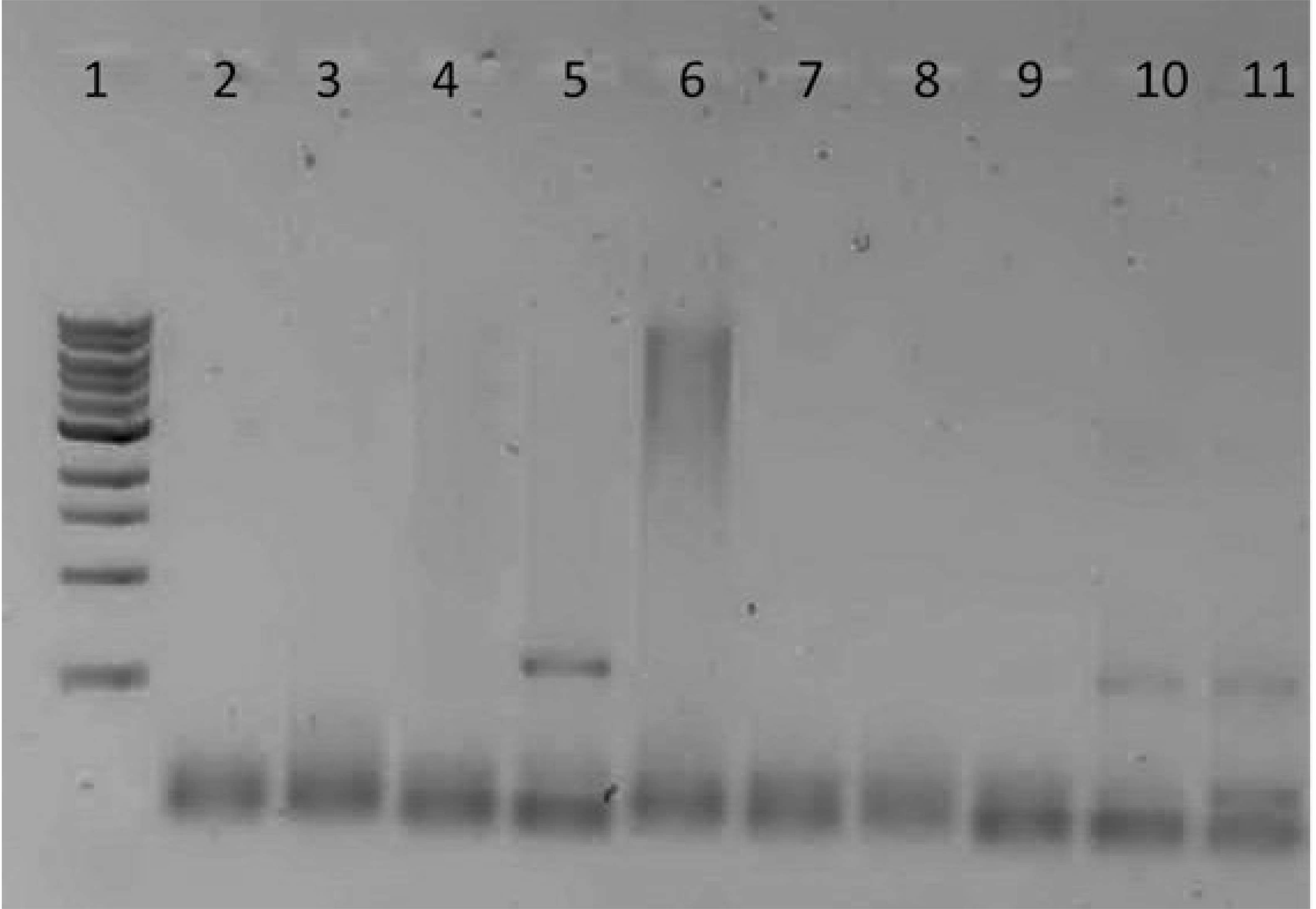
Agarose gels with putative xylanase gene fragment amplicons. The amplicons were obtained by Taq polymerase PCR using primer pairs specific for xylanases of glycohydrolase families 8 (A; xyn8A), 10 (B; xynFA), and 11 (C; PAxynA). All gels show a 1kb DNA ladder (NEB) in the first lane, showing bands from 0.5kb to 10kb. Lanes 2 to 11 contain amplicons from strains as follows: 2, AIBO216B; 3, AIBO216C; 4, AIBO228; 5, AL1515; 6, AL1610; 7, AL1614; 8, DO1702; 9, OV2120; 10, OV2121; and 11, OV2122.

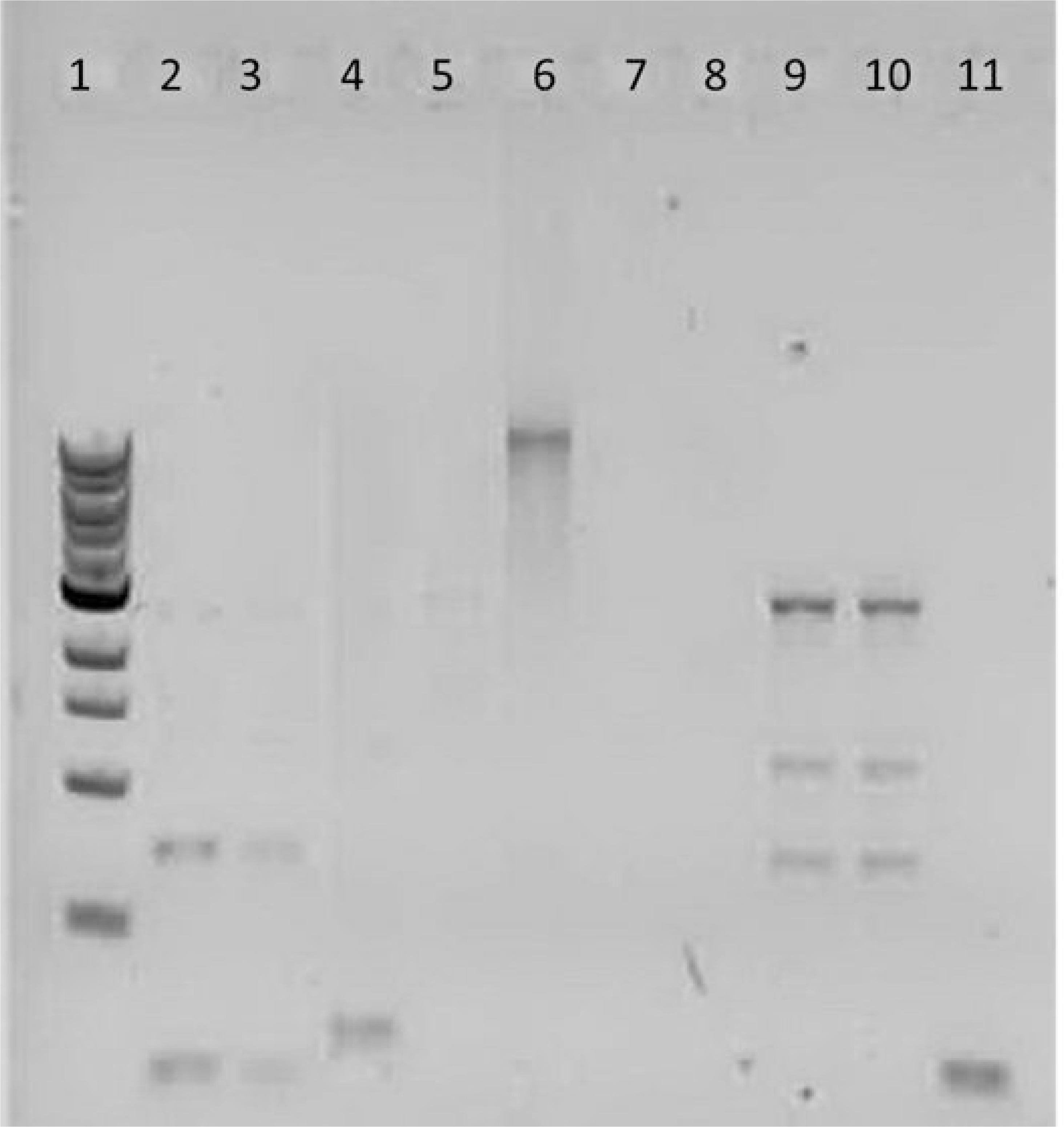

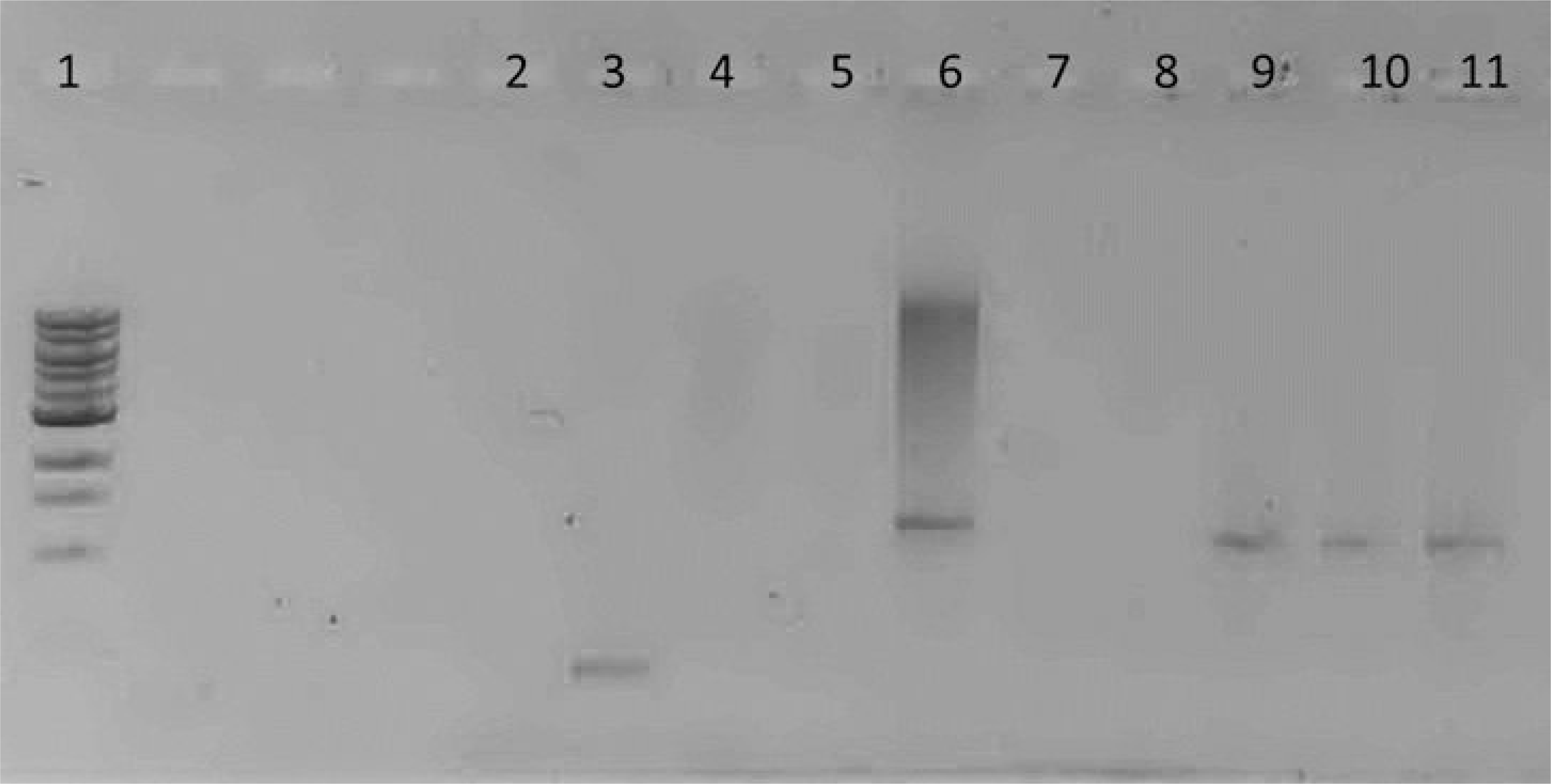

### Temperature-dependence of xylanolytic activity

The xylanase screen was adapted to a semi-quantitative assay by monitoring the emergence and diameter of the blue dye haloes on triplicate plates incubated at 5, 15, 22, and 35°C over a period of up to 10 days. The effect of incubation temperature on xylanolytic activity varied substantially among the strains (Figure 3), with OV2121 displaying the most prominent activity at 5°C, whereas at 35°C AL1614 and AL1515 displayed the most activity. Although variation among replicates was in many cases high (Table 6), especially at lower temperatures, one-way ANOVA on terminal halo diameters revealed significant differences in xylanase activity among the strains at 35°C (F_(7,16)_ = 8.45; p<0.001) and at 22°C (F_(8,16)_ = 4.45; p=0.005), while at 15°C one-way ANOVA did not reveal a difference (F_(7,16)_ = 1.44; p=0.257) and at 5°C, while evidence against the null hypothesis of there being no difference was found, it was not significant at the 95% level (F_(5,12)_ = 3.01; p=0.055). Most of the isolates displayed higher xylanolytic activity at higher temperatures, although isolate OV2122 only displayed activity below 35°C. No activity was detected at 5°C in any of the replicates of isolates AL1515, AL1610, or AL1614.

**Table 6.** Terminal halo diameters on Az-xylan test media at different incubation temperatures.

| *Strains* | 5°C | 15°C | 22°C | 35°C |
| --- | --- | --- | --- | --- |
| AIBO216B | 1.3 ± 0.4 | 3.5 ± 0.5 | 4.0 ± 0.0 | 4.0 ± 0.0 |
| AIBO216B | 0.7 ± 0.6 | 3.2 ± 0.3 | 4.2 ± 0.3 | 4.3 ± 0.3 |
| AIBO228 | 0.3 ± 0.6 | 2.5 ± 0.9 | 3.8 ± 0.3 | 3.8 ± 0.3 |
| AL1515 | 0.0 | 0.0 | 3.2 ± 0.3 | 6.3 ± 0.6 |
| AL1610 | 0.0 | 3.5 ± 0.7 | 3.6 ± 0.1 | 5.2 ± 0.3 |
| AL1614 | 0.0 | 2.0 ± 0.4 | 3.6 ± 0.8 | 6.5 ± 1.3 |
| OV2120 | 0.5 ± 0.5 | 3.3 ± 0.6 | 4.3 ± 0.4 | 4.3 ± 1.0 |
| OV2121 | 2.3 ± 1.6 | 3.5 ± 0.0 | 3.4 ± 0.1 | 3.7 ± 0.3 |
| OV2122 | 0.2 ± 0.3 | 2.3 ± 0.3 | 3.0 ± 0.5 | 0.0 |

**Figure 3.**
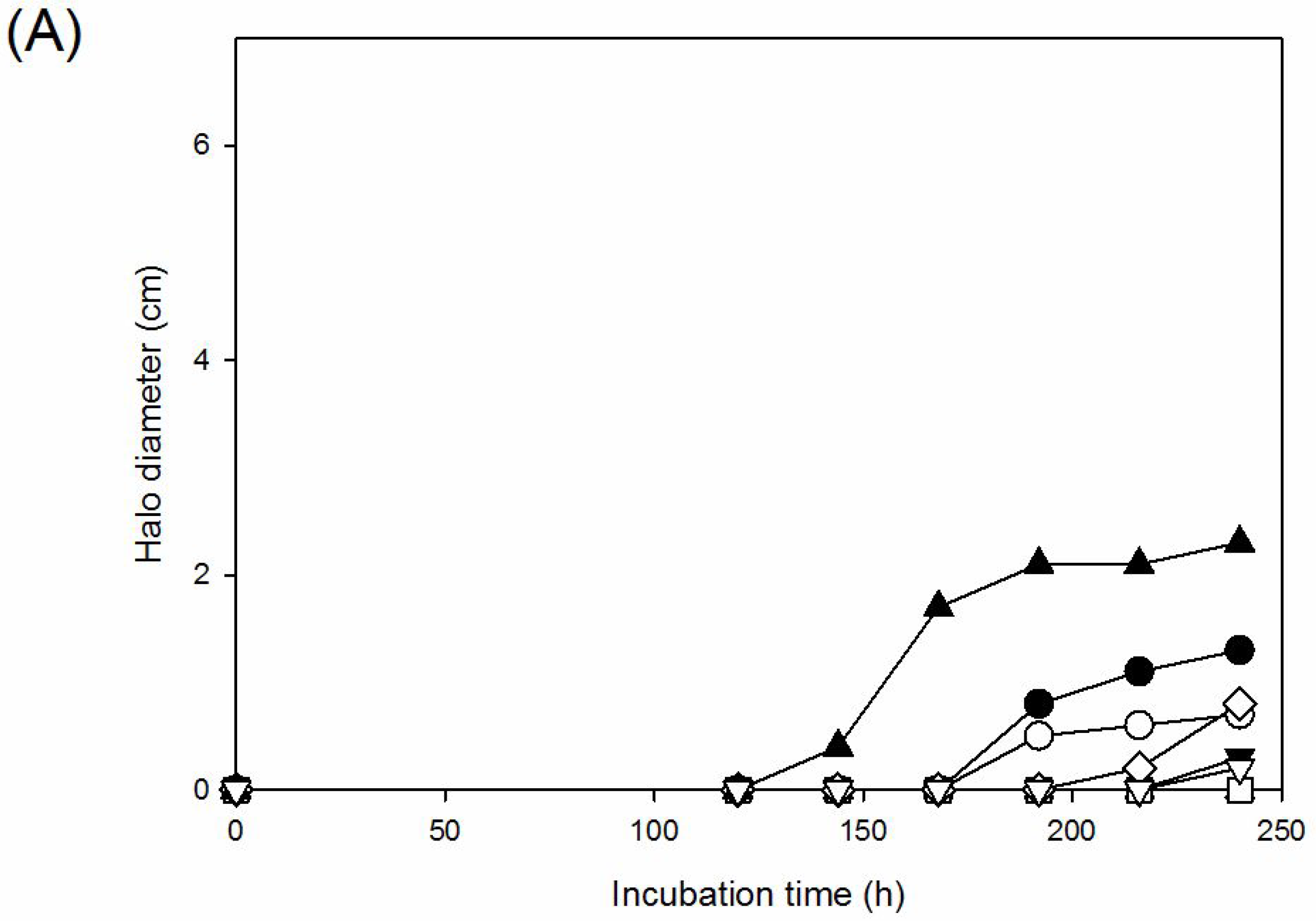
Xylanolytic activity as measured by the emergence and diameter of dye halos on azo-crosslinked birch xylan (Megazyme) agar media. Plates were incubated at 5°C (A), 15°C (B), 22°C (C), and 35°C (D). Values are means of measurements of triplicate plates. Error bars are omitted for clarity (standard deviations of terminal values can be seen in Table 6). Solid circles indicate strain AIB0216B, open circles strain AIB0216C, solid downwards-pointing triangles strain AIB0228, open upwards-pointing triangles strain AL1515, solid squares strain AL1610, open squares strains AL1614, open diamonds strain OV2120, solid upwards-pointing triangles strain OV2121, open downwards-pointing triangles strain OV2122.

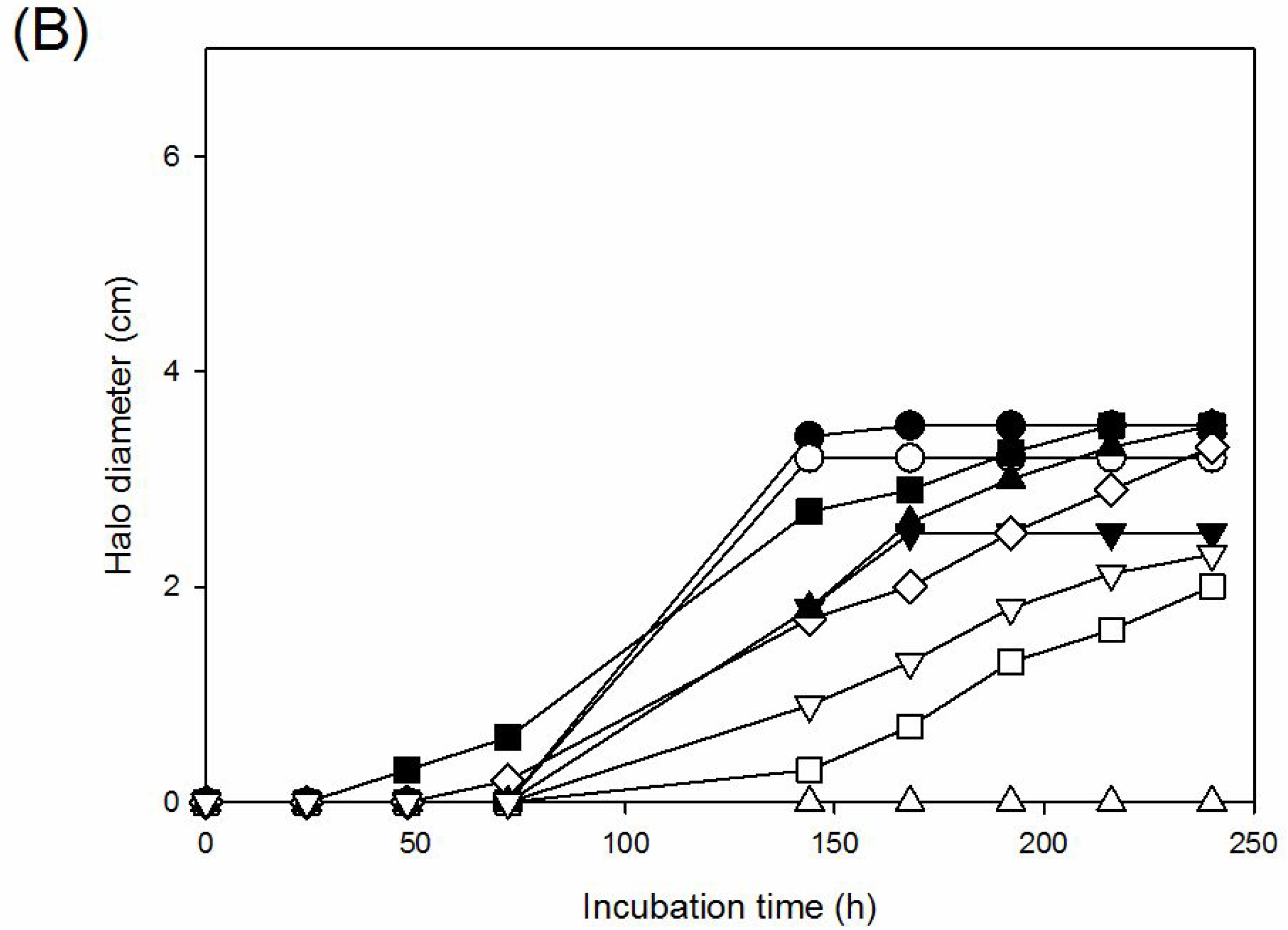

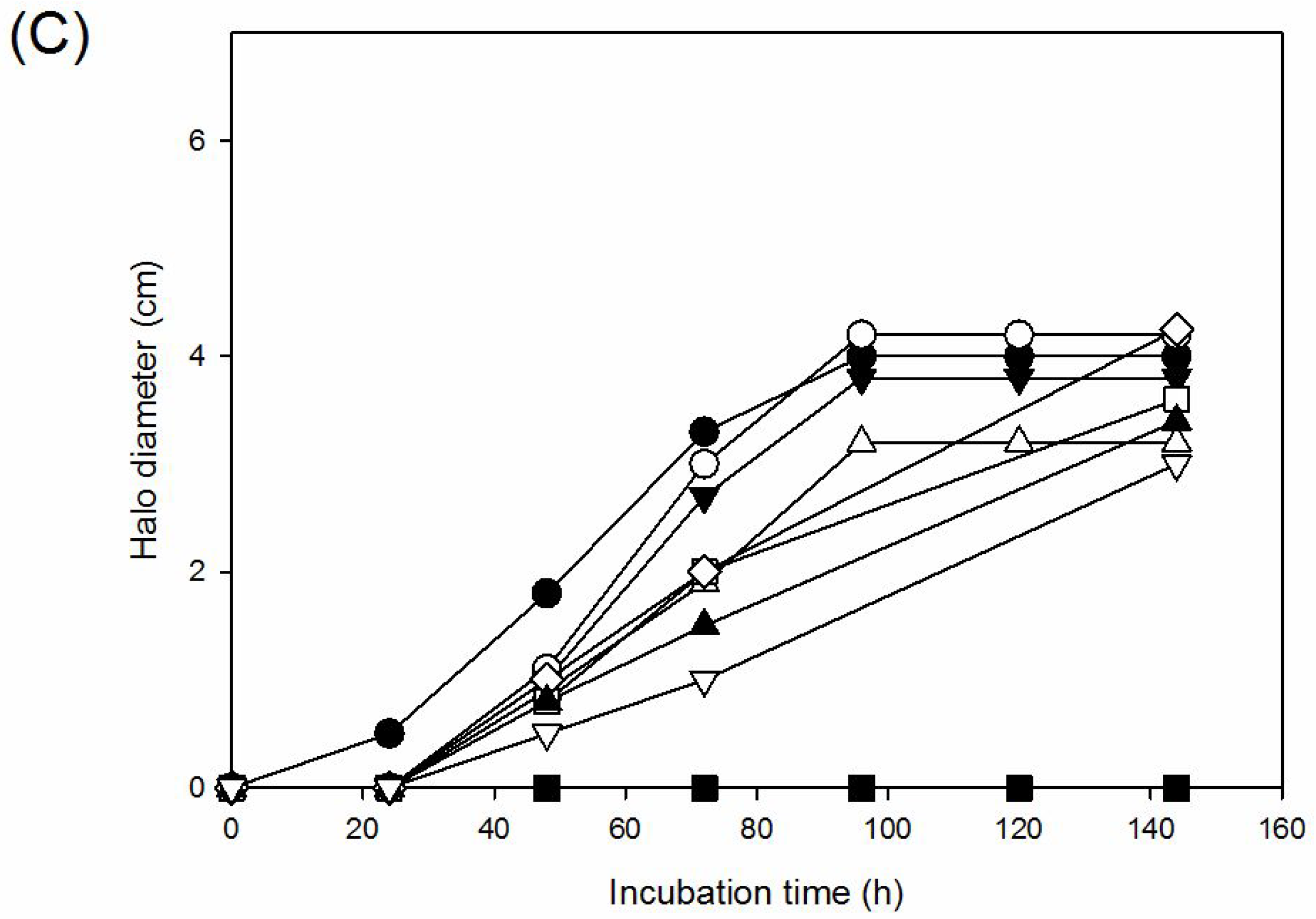

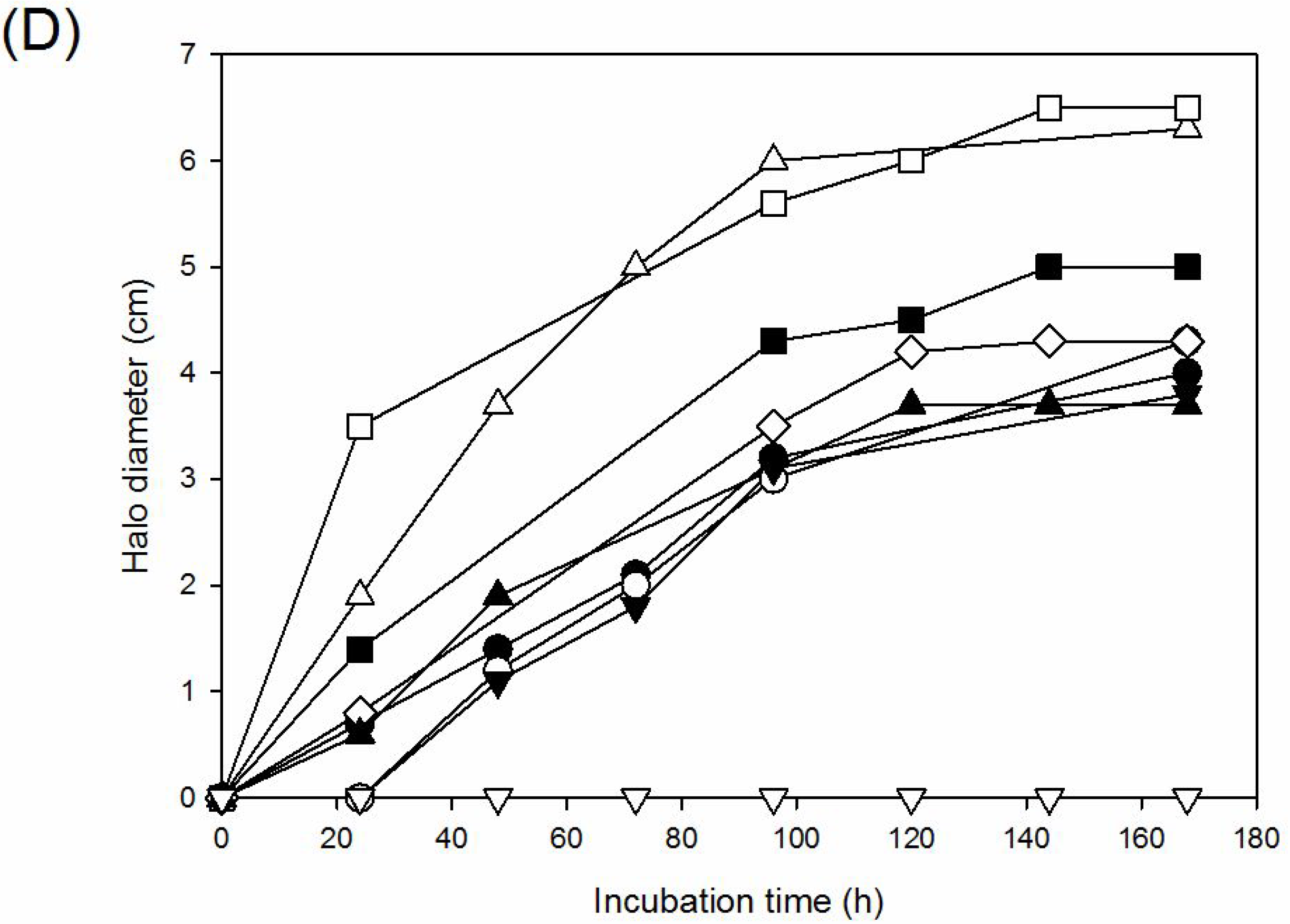

### Discussions

According to the 16S rRNA gene sequencing, five of the strains in present study belong to the genus of *Paenibacillus*. The paenibacilli are ubiquitous in nature, having been isolated from environments as diverse as soil, clinical specimens, and hot springs (16). Members of the genus are highly diverse, with member species representing psychrophiles, mesophiles, and thermophiles, strict aerobes and facultative anaerobes, and are known for their production of various glycanolytic exoenzymes (28). Since its formation from members of *Bacillus* (2), the genus *Paenibacillus* currently has grown to 200 species and 4 subspecies according to the LPSN (27). Xylanolytic activity is common among the paenibacilli (6) and several investigators have reported the isolation and characterization of glycohydrolase family 8 (23), family 10 (17, 37) and family 11 (34, 38) xylanases from these organisms. Several paenibacilli are psychrotrophic and, hence, cold-active xylanases are found among them (40).

Much like the paenibacilli, the pseudomonads are very commonly encountered in plant-and soil-associated environments, and a number of studies have described xylanolytic activity in, and the isolation and characterization of xylanases from these bacteria (20, 29). The *P. fluorescens* family 10 xylanase A and family 11 xylanase E have been well studied (13).

The identity of xylanolytic isolate AL1610 as a *Stenotrophomonas* sp. was more surprising. While the stenotrophomonads are typically plant-associated bacteria and therefore expected in the habitats under study, they are generally, as the genus name implies, quite fastidious with a nutritional spectrum mostly limited to mono-and disaccharides (25). Very little literature currently exists on xylanolytic activity in these organisms. Indeed, according to Raj et al. (30), their study on an inducible, thermostable xylanase from *Stenotrophomonas maltophilia* was the first to describe xylanase activity in that species. Malfliet et al. (21) also reported *S. maltophilia* xylanase activity among the consortia present in malting barley. The stenotrophomonads have been used in agricultural and environmental biotechnology, such as for bioremediation, in spite of the potential of *S. maltophilia* as an opportunistic human pathogen (4). Xylanolytic activity may further expand the biotechnological potential of these organisms. A pairwise distance comparison of the aligned 16S rRNA sequences, obtained for all nine strains, revealed a close relatedness of strains isolated in widely separated parts of the country. These results are striking but not necessarily unexpected, as the sporeforming paenibacilli disperse easily by wind over considerable distances and are thus not subjected to the physical isolation required for diversification (26). The fifth *Paenibacillus*, strain OV2122 was more divergent, displaying on average 67 base substitutions to each of the other paenibacilli over the 987 nucleotide positions tested. The other four strains were more heterogeneous, although the two pseudomonads were fairly similar to each other (9 substitution over the 987 positions). The highest divergence observed was between *Paenibacillus* OV2122 and *Stenotrophomonas* AL1610, 298 base substitutions among the 987 nucleotide positions tested. Four isolates, OV2121, OV2120, AIB0216B and AIB0216C, form a clade with *P. amylolyticus*, *P. xylanexedens*, *P. tundae* and *P. tylopili* (Figure 1). Both *P. xylanexedens* and *P. tundrae* were originally isolated as xylanolytic paenibacilli from moist tundra soil in Alaska (24). However, related strains have been isolated from other environments, such as barley grain in Finland (31) and catfish guts in Brazil (8). Isolate OV2122 forms a clade with *P. castaneae* and *P. endophyticus* supporting the EzTaxon assignment of this isolate as *P. castaneae*, a species described from strains originally isolated from a chestnut tree rhizosphere in Spain (39).

Several of the strains yielded amplicons from more than one xylanase primer pair used for screening. While this may be due to less-than-perfect specificity of the primers employed due to the noted diversity of xylanase amino acid sequences (7), more than one xylanase genes may indeed be present in the respective genomes. Several microorganisms, including both pseudomonads and bacilli, are known to produce multiple xylanases (11, 41). Taken together, these results indicate that the paenibacilli (strains AIB0216B, AIB216C, OV2120, OV2121, and OV2122) possess at least family 10 and family 11 xylanases, strains OV2121 and OV2122 possibly also a family 8 xylanase. The two pseudomonads (AIB0228 and AL1515) display quite divergent results, with AL1515 possibly possessing a family 8 xylanase, although in both cases the identity of the PCR amplicons could not be verified by sequencing as belonging to a xylanase gene. Based on the 261-bp PAxynA amplicon sequence, the *Stenotrophomonas* AL1610 xylanase appears most similar to the family 11 xylanase of *Paenibacillus* HY-8 (14).

We adapted the xylanase screen to a semi-quantitative assay and monitored the emergence and diameter of the blue dye haloes incubated at different temperature, ranging from 5 – 35°C. The effect of temperature was noticeable most of the tested strains displayed most activity at higher temperature. Of the nine tested strains, only *Paenibacillus amylolyticus* OV2121, displayed the most prominent activity at 5°C. We therefore conclude that of the nine xylanolytic strains studied, *Paenibacillus amylolyticus* OV2121 produces the most cold-active xylanase under the conditions employed, while *Pseudomonas kilonensis* AL1515 and *Bacillus subtilis* AL1614 display higher activity at higher temperatures. *Paenibacillus castaneae* OV2122 only displays xylanolytic activity at lower temperatures.

## Acknowledgments

This work was in part conducted under an ERASMUS internship exchange programme and was funded by the University of Akureyri Research Fund. The authors would also like to acknowledge Robert W. Jackson and other coordinators of the University of Akureyri and Reading University joint Arctic microbiology field trip and laboratory course of 2012 within which environmental sampling and initial culturing of the isolates was performed by students Árný Ingveldur Brynjarsdóttir and Luke Johnson.

